# Chronic sensing of host-derived lipids is an all-in-one signal that primes and activates NLRP3

**DOI:** 10.1101/2025.10.29.685328

**Authors:** Océane Dufies, Marco Di Gioia, Roberto Spreafico, Piao J Tan, James R Springstead, Sean A Prell, Junning Case, Ce Gao, Kevin Wei, Hao Wu, Ivan Zanoni

**Author notes:** Corresponding author: Ivan Zanoni.

## Abstract

Activation of the NLRP3 inflammasome leads to the production of bioactive interleukin (IL)-1β fostering atherosclerosis. The current dogma is that NLRP3 must be first primed by microbial stimuli, known as pathogen-associated molecular patterns (PAMPs), and then activated by either microbial or host-derived inflammatory cues. The mechanism that controls NLRP3 functioning in the context of non-communicable diseases lacking overt microbial infections remains debated. Here, we show that chronic exposure to atherosclerosis-associated oxidized phospholipids (oxPLs) simultaneously primes and activates NLRP3 independently of microbial cues. Mechanistically, chronic exposure to host-derived oxPLs activate the transcription factor NRF2, which is necessary and sufficient to prime and activate NLRP3 in a PAMP-independent manner. NRF2 chronic activation drives oxidized mitochondrial DNA to activate NLRP3. *Ex vivo* analyses of atherosclerotic plaques in mice and humans identify a population of monocytes-derived macrophages which activates NRF2 and expresses IL-1β. Overall, our data point to oxPL-dependent NRF2 activation as an all-in-one signal necessary and sufficient to prime and activate NLRP3, sustaining atherogenesis.

## Introduction

Metabolic inflammation, or metaflammation, is a major driver of non-communicable inflammatory diseases^1^. Metaflammation sustains the development of cardiovascular diseases and atherosclerosis, which represent a major cause of death worldwide^2, 3^. Inflammasomes are supramolecular complexes (SMOCs) that drive the activation of inflammatory caspases, cleaving multiple substrates and ultimately leading to the secretion of bioactive IL-1β^4^, that is a major therapeutic target in atherosclerotic patients^3, 5^. The NLRP3 inflammasome is the key mediator of IL-1β production during atherosclerosis^6^, and its activity strictly relies on a two-step process that requires a priming phase followed by an activation step^7-10^. The priming step of NLRP3 is usually driven by detection of pathogen-associated molecular patterns (PAMPs) by pattern recognition receptors (PRRs). PRR activation provides the priming signal via NF-κB-dependent upregulation of pro-IL-1β and NLRP3, and via post-translational modifications of the NLRP3 SMOC^7, 11, 12^. Following priming, microbial- or host-derived inflammatory cues drive NLRP3 activation. Although a variety of signals can activate NLRP3, they often converge onto mitochondrial ROS production and cytosolic accumulation of oxidized mitochondrial DNA which binds and activates NLRP3^13-16^. It remains a mystery whether host-derived damage-associated molecular patterns (DAMPs) produced during atherogenesis in the absence of an overt infection can simultaneously prime and activate the NLRP3 inflammasome, thus sustaining metabolic diseases.

Atherosclerotic patients are characterized by an elevated concentration of low-density lipoprotein (LDL) cholesterol and the accumulation of oxidized LDL (oxLDL) in the intima of arteries. Following endocytosis, soluble oxLDL nucleates into particulates that activate the NLRP3 inflammasome^17-19^. A set of heterogenous oxidized phospholipids (oxPLs), collectively known as oxidized 1-palmitoyl-2-arachidonoyl-*sn*-phosphatidylcholine (oxPAPC), is recognized as the bioactive moiety that drives inflammation in response to oxLDL encounter^20^. A growing body of evidence supports the pathological role of the oxPAPC in metabolic diseases^21-25^, and distinct components of oxPAPC (such as PGPC and POVPC) have been shown to activate the NLRP3 inflammasome in the context of microbial encounter^26-29^. Curiously, the acute administration of oxPAPC fails to induce IL-1β secretion in PAMP-primed macrophages^26, 27^, raising questions about the relevance of this DAMP in regulating inflammasome activation in general, and during atherosclerosis development in particular.

To solve the conundrum of how the activity of the NLRP3 inflammasome is sustained in a non-communicable metabolic disease such as atherosclerosis, we tested the hypothesis that the chronic exposure to oxPAPC controls NLRP3 inflammasome activation and we assessed how this process impacts atherosclerosis development.

## Results

### Chronic exposure to oxPAPC is sufficient to license the activity of the NLRP3 inflammasome

Macrophages primed with microbial ligands and then acutely treated with oxPAPC do not activate the inflammasome^26, 27^. Since atherosclerotic patients are chronically exposed to high levels of oxPLs in their blood^20^, we sought to test whether pre-conditioning of myeloid cells with oxPLs affects their capacity to activate the inflammasome. Macrophages were either treated with oxPAPC for 24 hours before stimulation with Toll-like receptor (TLR) agonists, or, as a control, they were acutely treated with oxPAPC after priming of cells with PAMPs, and IL-1β secretion was used as a proxy of inflammasome activation (**Figure 1A**). Conditioning of macrophages with oxPAPC for 24 hours followed by TLR stimulation led to IL-1β secretion in a dose-dependent manner (**Figures 1B and 1C**). In contrast, and in keeping with our previous results^26, 27^, administration of the whole oxPAPC mixture after TLR stimulation did not induce IL-1β secretion (**Figure 1B**). Of note, administration of PGPC, that activates the non-canonical inflammasome pathway^26-29^, induced IL-1β secretion only when administered after, but not prior to, TLR stimulation (**Figure 1B**). These data suggest that the pre-exposure of macrophages to the mixture of oxPLs contained in oxPAPC licenses inflammasome activation via a different process than the one previously described for PGPC. In keeping with this hypothesis, caspase-11 and CD14, that are required for PGPC-mediated inflammasome activation^26-29^, were dispensable for IL-1β secretion by macrophages conditioned with oxPAPC before TLR stimulation (**Figures S1A and S1B**).

**Figure 1:**
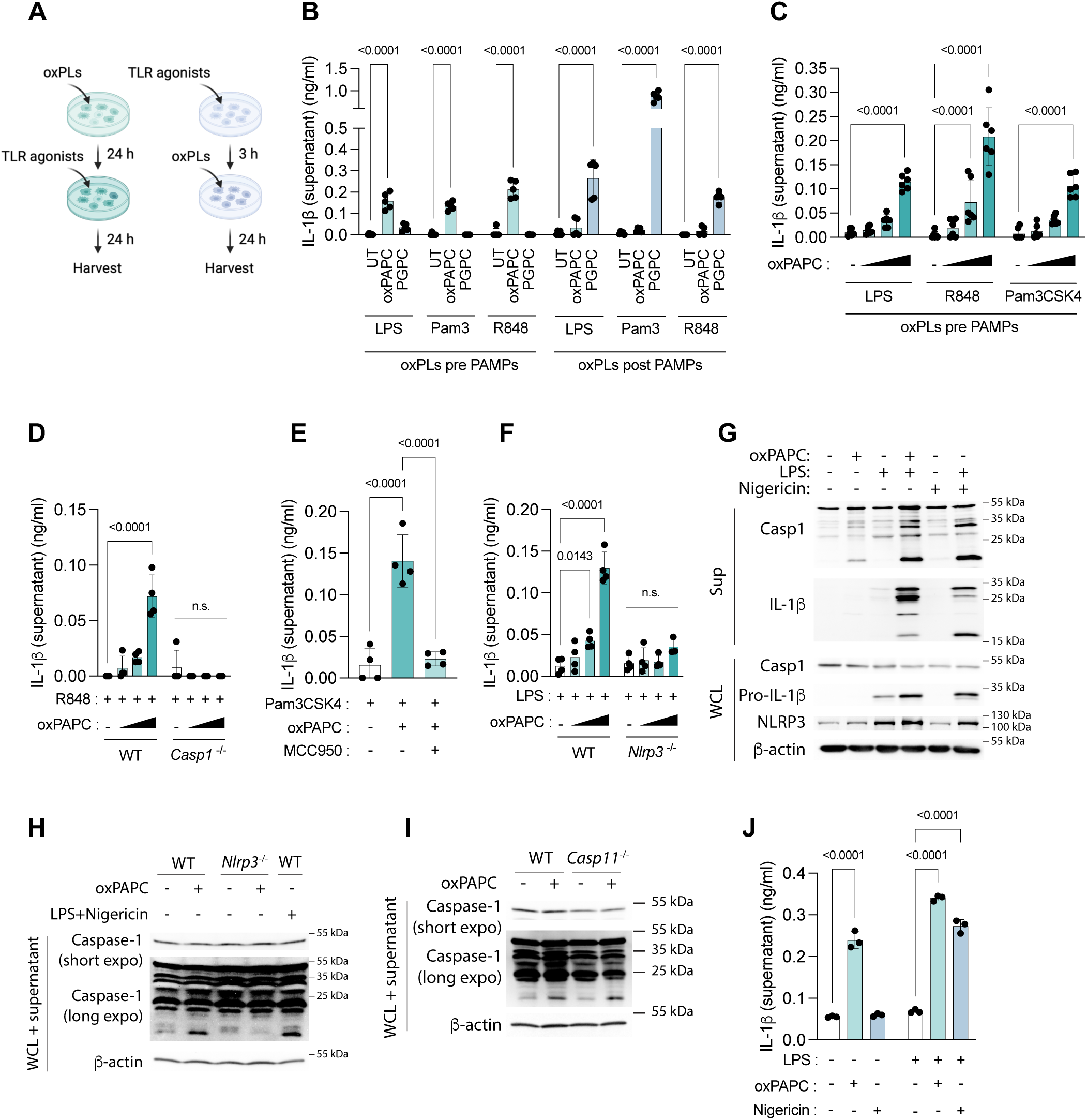
Chronic exposure to oxPAPC is sufficient to license the activity of the NLRP3 inflammasome. (**A**) Experimental design. Bone marrow-derived macrophages (BMDMs) were either conditioned with oxPLs (oxPAPC or PGPC) for 24 hours (h) and treated with TLRs agonists (Pam3CSK4, R848 or LPS) for 24 h (*left*), or primed for 3 hours with TLR agonists and treated with oxPLs for 24 h (*right*). Made with BioRender. (**B**) BMDMs were treated as indicated in (**A**) and IL-1β in the supernatant was assessed by ELISA. (**C, D**) WT (**C, D**) or *Casp1*^-/-^ (**D**) BMDMs were incubated with oxPAPC (25, 50 and 100 µg/ml) for 24 h and treated with TLR agonists (1 µg/ml) as indicated (**C**) or with R848 for 24 h (**D**). IL-1β in the supernatant was assessed by ELISA. (**E**) BMDMs were pre-treated for 1 h with MCC950 (5 µM) before conditioning for 24h with oxPAPC (100 µg/ml). Cells were treated for 24 h with Pam3CSK4 (1 µg/ml). Cells were analyzed as in (**C**). (**F**) WT or *Nlrp3*^-/-^ BMDMs were conditioned for 24 h with oxPAPC (25, 50 and 100 µg/ml) and treated for 24 h with LPS (1 µg/ml). Supernatants were analyzed as in (**C**). (**G**-**I**) WT (**G**-**I**), *Nlrp3*^-/-^ (**H**), or *Casp11*^-/-^ (**I**) BMDMs were either conditioned with oxPAPC (100 µg/ml) for 24 h and treated, or not, with LPS (1 µg/ml), or treated with Nigericin (5 µM) alone or primed for 4 h with LPS (1 µg/ml) and then stimulated with Nigericin for 1 h, as indicated. Supernatants (Sup) and whole cell lysates (WCL) were analyzed by immunoblot. (**J**) THP-1 monocytes expressing mTurquoise2-hProIl1b-mNeonGreen were treated as in (**G**) and IL-1β in the supernatant was assessed by ELISA. Graphs show means ± SD and statistical significance was calculated using a Two-way ANOVA.

Next, we tested whether IL-1β secretion by macrophages pre-conditioned with oxPAPC was dependent on NLRP3 inflammasome and caspase-1 activation. We utilized macrophages that were either deficient for *Casp1* or *Nlrp3*, or that were treated with the NLRP3 inhibitor MCC950. Exposure of cells to TLR agonists demonstrated that both caspase-1 and NLRP3 are required to trigger IL-1β secretion in macrophages pre-conditioned with oxPAPC (**Figures 1D** – **F**). In contrast, levels of pro-IL-1β or IL-6 were not affected in the presence or absence of NLRP3 (**Figures S1C and S1D**). Similar to acute exposure to oxPAPC^26-29^, but in contrast to cells treated with a PAMP and a canonical NLRP3 activator such as ATP, cells pre-conditioned with oxPAPC were hyperactive^26-29^ and did not encounter pyroptotic cell death (**Figure S1E**).

Our data so far demonstrated that conditioning of macrophages with oxPAPC licenses NLRP3-dependent IL-1β secretion upon TLR stimulation. Under these experimental conditions, TLR signaling may drive the PTMs required for NLRP3 priming, or it could simply increase the levels of pro-IL-1β for subsequent cleavage by a primed and activated inflammasome. To discriminate these two possibilities, we performed immunoblotting to assess inflammasome activity by measuring the cleavage not only of IL-1β, but also of caspase-1, that does not require to be upregulated following PAMP encounter. Lipopolysaccharide (LPS) administration, but not oxPAPC only, induced pro-IL-1β accumulation (**Figure 1G**). TLR stimulation of macrophages pre-conditioned with oxPAPC triggered the cleavage of both pro-caspase-1 and pro-IL-1β into active caspase-1 (20 kDa) and mature IL-1β (17 kDa), as observed when NLRP3 was activated with LPS and Nigericin, another canonical activator of the NLRP3 inflammasome (**Figure 1G**). Surprisingly, pre-treatment of macrophages with oxPAPC in the absence of TLR stimulation was sufficient to induce the cleavage of caspase-1, while, as expected, Nigericin failed to do so (**Figures 1G** – **I**). Caspase-1 activation in response to oxPAPC conditioning was abolished in *Nlrp3*^-/-^, but not *Casp11*^-/-^, macrophages (**Figures 1H and 1I**).

Finally, we reasoned that conditioning with oxPAPC could trigger IL-1β maturation and secretion in cells that spontaneously express high levels of pro-IL-1β. To test this hypothesis, we took advantage of a human monocytic cell line that constitutively express high pro-IL-1β^30^. When cells were treated with Nigericin, they secreted IL-1β only if they were previously primed with LPS (**Figure 1J**). In contrast, exposure to oxPAPC for 24 hours, either in the absence or presence of TLR stimulation, led to secretion of IL-1β, in a NLRP3-dependent manner (**Figures 1J and S1F**).

Overall, these data indicate that chronic exposure to oxPAPC is sufficient to drive NLRP3 inflammasome activation in the absence of PAMPs.

### oxPAPC-dependent activation of NRF2 is necessary and sufficient to drive the activity of the NLRP3 inflammasome

Next, we tested the duration of oxPAPC pre-conditioning necessary to promote IL-1β secretion by macrophages. We found that cells required to be exposed to oxPAPC at least for 4-8 hours to acquire the capacity to secrete IL-1β, while IL-1β secretion peaked when cells were pre-conditioned with oxPAPC for 24 hours (**Figure 2A**). This kinetic led us to hypothesize that oxPAPC induces transcriptional changes necessary for licensing the activity of NLRP3. To explore this possibility, we performed bulk RNA sequencing (RNAseq) on macrophages treated for 3 or 24 hours with oxPAPC. oxPAPC reprogrammed gene expression of macrophages both at 3 and 24 hours (**Figures 2B and 2C**). Ingenuity Pathway Analysis (IPA) and gene set enrichment analysis (GSEA) revealed that genes regulated by the transcription factor NRF2 were highly upregulated by oxPAPC at all time points (**Figures 2D**, **S2A**). oxPAPC administration to macrophages induced NRF2 stabilization and nuclear translocation as early as 2 hours and up to 24 hours (**Figures 2E and S2B – D**).

**Figure 2:**
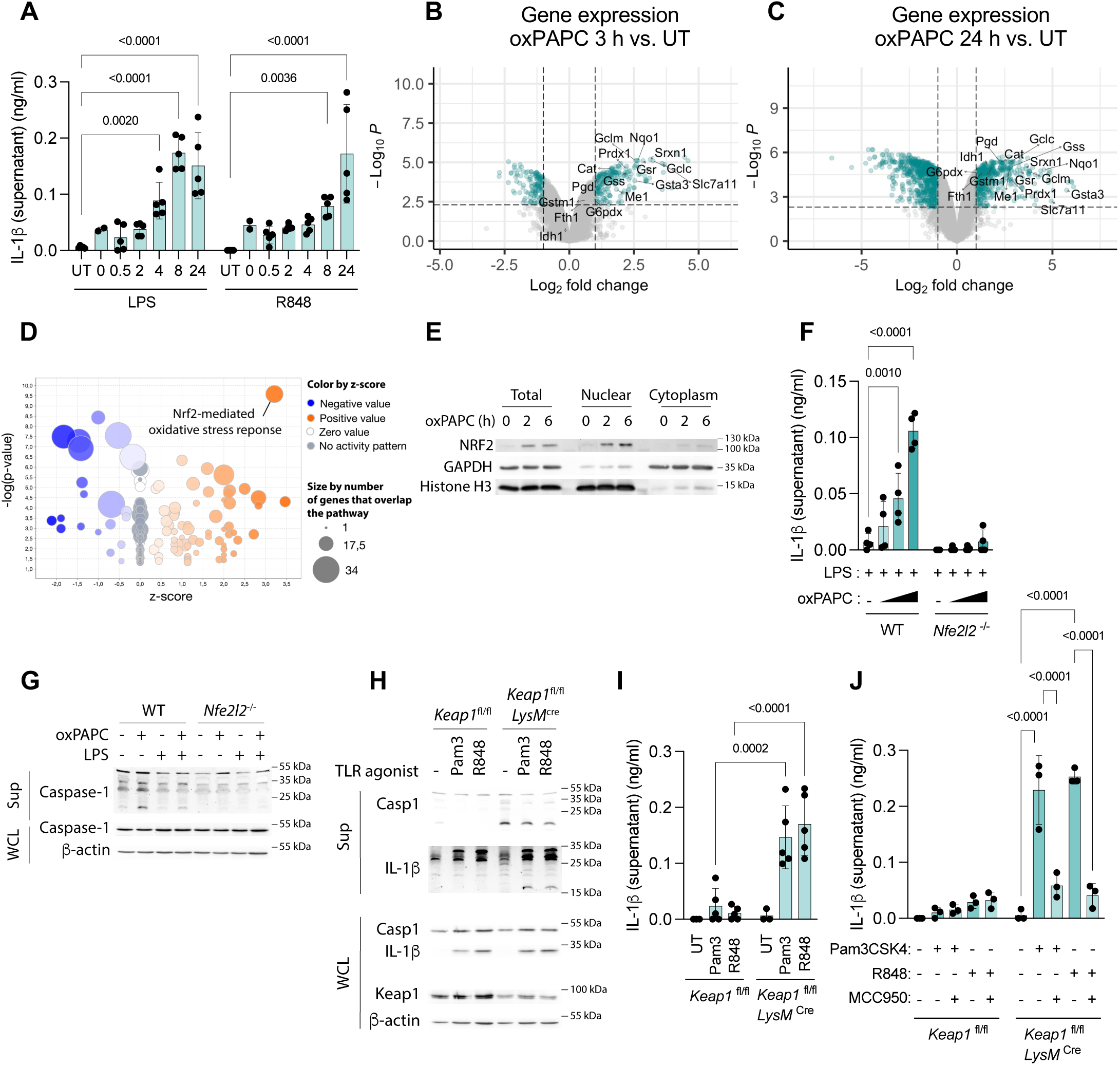
NRF2 activation is necessary and sufficient to prime and activate the NLRP3 inflammasome in response to chronic exposure to oxPAPC. (**A**) BMDMs were preconditioned, or not (UT), with oxPAPC (100 µg/ml) for the indicated time (0 to 24 h) before adding LPS or R848 (1 µg/ml) for 24 h. IL-1β in the supernatant was assessed by ELISA. (**B**-**D**) BMDMs were incubated, or not (untreated, UT), with oxPAPC (100 µg/ml) for 3 h (**B**) or 24 h (**C, D**) before being analyzed by bulk RNAseq. (**B, C**) Volcano plot indicating the genes up- and down-regulated by oxPAPC at 3 h (**B**) and 24 h (**C**). Dotted lines and turquoise coloring represent fold change > 2 and p-value < 0.01. Major NRF2-regulated genes are highlighted. (**D**) Bubble plot of Ingenuity Pathway Analysis of canonical pathways induced by oxPAPC conditioning for 24 h. (**E**) BMDM were treated for the indicated time with oxPAPC (100 µg/ml) and cells were subjected to nuclear-cytoplasmic fractionation and analyzed by immunoblot. (**F, G**) WT or *Nfe2l2*^-/-^ BMDMs were pre-conditioned with oxPAPC (100 µg/ml) for 24 h before adding, or not, LPS for 24 h (1 µg/ml), as indicated. IL-1β in the supernatant was assessed by ELISA (**F**). Supernatants (Sup) and whole cell lysates (WCL) were analyzed by immunoblot (**G**). (**H**-**J**) *Keap1*^fl/fl^ or *Keap1*^fl/fl^*LysM*^cre^ BMDMs were pre-treated (**J**), or not (**H, I**), with MCC950 (5 µM) for 1 h before treatment with Pam3CSK4 (Pam3) or R848 (1 µg/ml) for 24 h. Supernatants and whole cell lysates were analyzed by immunoblot (**H**). Supernatants were analyzed by ELISA (**I, J**). Bar graphs show means ± SD and statistical significance was calculated using a Two-way ANOVA. Volcano plots show differential expression and adjusted p-value using *limma*.

NRF2 has been widely reported as a negative regulator of NLRP3 activation^31-37^, although scattered reports suggested NRF2 can also positively regulate the NLRP3 inflammasome^38-40^. To assess the role of NRF2 in our experimental system, we utilized *Nfe2l2*^-/-^ macrophages that lack NRF2 (**Figure S2B**) and tested whether NRF2 was necessary to activate NLRP3 in response to oxPAPC. *Nfe2l2*-deficency abrogated the capacity of macrophages pre-conditioned with oxPAPC to secrete IL-1β (**Figure 2F**). Caspase-1 activation was abrogated in *Nfe2l2*^-/-^ cells treated with oxPAPC, either in the presence or absence of LPS (**Figure 2G**), while the levels of pro-IL-1β or IL-6 were only marginally affected in wild-type (WT), compared to *Nfe2l2*^-/-^, macrophages (**Figures S2E and S2F**). Moreover, under our experimental conditions, the secretion of IL-1β induced by canonical NLRP3 activators was not affected in *Nfe2l2*^-/-^ macrophages (**Figure S2G**), as also previously reported^41^.

To investigate whether NRF2 activation is not only necessary, but also sufficient to induce the activity of the NLRP3 inflammasome, we used *Keap1*^fl/fl^*LysM*^cre^ macrophages that bear a constitutively active NRF2^42^, as indicated by the upregulation of NRF2-dependent genes (**Figure S2H**). These cells, compared to *Keap1*^fl/fl^ controls, showed activated caspase-1 in the absence of any stimulation and, upon TLR-dependent induction of pro-IL-1β (**Figure S2I**), spontaneously secreted bioactive IL-1β in a NLRP3-dependent manner (**Figures 2H** – **J**). No major changes in the levels of secreted IL-1β were detected when *Keap1*^fl/fl^*LysM*^cre^ or *Keap1*^fl/fl^ macrophages were treated with canonical activators of NLRP3 (**Figure S2J**). Potassium efflux is a major driver of inflammasome activation in response to canonical NLRP3 activators^43, 44^, and, indeed, potassium efflux blockade prevented IL-1β secretion in response to Nigericin (**Figure S2K**). In contrast to Nigericin, cells pre-conditioned with oxPAPC or that lack KEAP1 secreted IL-1β independently of potassium efflux (**Figures S2L and S2M**).

Altogether, these results demonstrate that NRF2 activation is necessary and sufficient to elicit the activity of NLRP3 in response to chronic exposure to oxPAPC.

### oxPAPC activates the NLRP3 inflammasome via NRF2-mediated accumulation of oxidized mitochondrial DNA

We, next, sought to further investigate how oxPAPC-dependent NRF2 signaling control the activation of NLRP3. The cytosolic export of oxidized mitochondrial DNA (ox-mtDNA) formed in response to mitochondrial reactive oxygen species (mtROS) activates NLRP3^14-16^. We, thus, evaluated whether oxPAPC drives mtROS. Treatment of macrophages with oxPAPC led to mtROS production (**Figure 3A**). *Keap1*^fl/fl^*LysM*^cre^ macrophages also displayed increased mtROS compared to their *Keap1*^fl/fl^ counterpart under homeostasis (**Figure 3B**). Quenching of mtROS reversed the capacity of cells conditioned with oxPAPC to activate caspase-1 (**Figure 3C**), or to secrete IL-1β (**Figure 3D**). mtROS production was accompanied by the accumulation of oxidized DNA both in oxPAPC-conditioned macrophages and *Keap1*-deficient cells (**Figures 3E – H**). Of note, NRF2 was required for the accumulation of oxidized DNA (**Figure 3I**). We, next, inhibited the endonuclease FEN1 as well as the mitochondrial channel VDAC1, that respectively processes ox-mtDNA and allows its cytosolic export to drive NLRP3 activation^15^. Blockade of VDAC1 or FEN1 significantly impaired the secretion of IL-1β in macrophages pre-exposed to oxPAPC (**Figure 3J**). Similar results were obtained upon inhibition of VDAC1 in *Keap1*^fl/fl^*LysM*^cre^ macrophages (**Figure 3K**). Overall, these data demonstrate that oxPAPC-mediated activation of NRF2 is responsible for mtROS production and release of ox-mtDNA into the cytosol, the latter serving as the activation signal for NLRP3.

**Figure 3:**
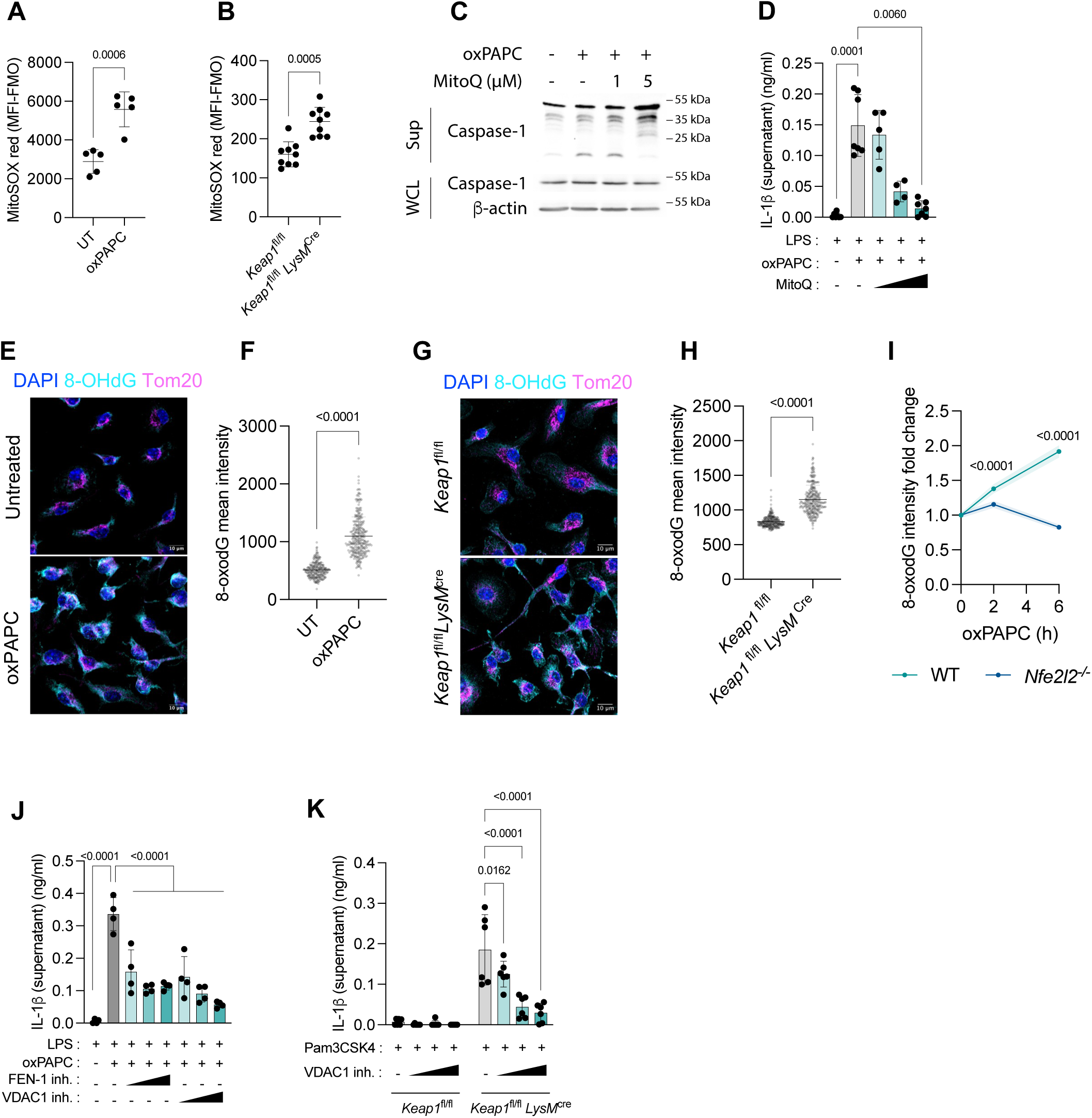
oxPAPC induces mitochondrial ROS to activate the NLRP3 inflammasome. (**A**) WT BMDMs were treated, or not, for 2 h with oxPAPC (100 µg/ml) and mitochondrial ROS were assessed with MitoSOX red. Cells were analyzed by flow cytometry. The median of fluorescence minus FMO (fluorescence minus one) is shown. (**B**) *Keap1*^fl/fl^ or *Keap1*^fl/fl^*LysM*^cre^ BMDMs were left untreated and analyzed as in (**A**). (**C, D**) WT BMDMs were pre-treated with mitoquinol (mitoQ, 1 µM, 2.5 µM or 5 µM) for 1 h before conditioning with oxPAPC (100 µg/ml) for 24 h and treated (**D**), or not (**C**), with LPS (1 µg/ml) for 24 h. Supernatants (Sup) and whole cell lysates (WCL) were analyzed by immunoblot (**C**). IL-1β in the supernatant was assessed by ELISA (**D**). (**E-H**) WT BMDMs were exposed to oxPAPC (100 µg/ml) for 24 h (**E, F**) and *Keap1*^fl/fl^ or *Keap1*^fl/fl^*LysM*^cre^ BMDMs were left untreated (**G, H**). Cells were stained by immunofluorescence for 8-OHdG (cyan, oxidized DNA), Tom20 (magenta, mitochondria) and DAPI (blue, nuclei) (**E, G**) and 8-OHdG intensity was quantified (**F, H**). Images show standard deviation projection of z-stack acquired with the Airyscan mode of Zeiss LSM880. Scale bar: 10 µm. (**I**) WT or *Nfel2l2*^-/-^ BMDM were treated for 0 to 6 hours with oxPAPC and stained for 8-OHdG as in (**E**). 8-OHdG intensity was measured and normalized to untreated condition. (**J, K**) WT (**J**), *Keap1*^fl/fl^ or *Keap1*^fl/fl^*LysM*^cre^ (**K**) BMDMs were pre-treated with a FEN1 inhibitor (FEN1-IN-4, 1 µM, 5 µM or 10 µM) (**J**) or a VDAC1 inhibitor (VBIT4, 1 µM, 5 µM or 10 µM) (**J, K**) for 1 h before conditioning (**J**), or not (**K**), with oxPAPC (100 µg/ml) for 24 h and treated with LPS (**J**) or Pam3CSK4 (**K**) (1 µg/ml) for 24 h. Supernatants were analyzed by ELISA. Bar graphs show means ± SD and statistical significance was calculated using a Two-way ANOVA (**D**, **J** and **K**) and t-tests (**A**, **B**, **F**, **H** and **I**).

Taken together, these results indicate that oxPAPC, via NRF2 activation, prime the NLRP3 inflammasome, allowing its activation in response to ox-mtDNA.

### NRF2 sustains atherogenesis and is activated in mouse and human monocyte-derived cells that respond to oxPAPC and produce IL-1β

Given that NRF2 controls the activity of NLRP3, we investigated the role of NRF2 in hematopoietic cells during the pathogenesis of atherosclerosis. *Ldlr*-deficient mice were reconstituted with WT or *Nfe2l2*-deficient bone marrow cells (**Figure 4A**) and fed a high-fat diet for 4 or 10 weeks. Levels of blood cholesterol were similar in the two groups (**Figure S3A**). The extent of atherosclerotic lesions and the formation of the necrotic core in the aortic roots were significantly decreased at early and late time points in mice that lacked NRF2 in hematopoietic cells, compared to mice reconstituted with WT bone marrow cells (**Figures 4B – E**), highlighting a detrimental role for NRF2 during atherogenesis.

**Figure 4:**
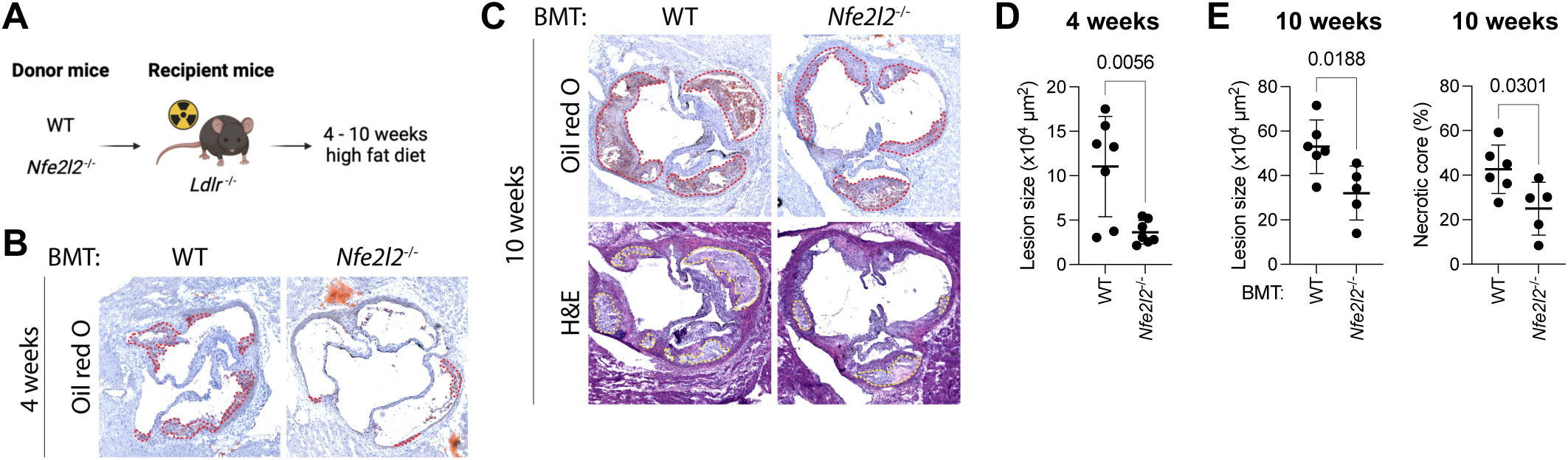
NRF2 activation in monocytes drives the formation of the atherosclerotic plaque. (**A**) Experimental design. *Ldlr*^-/-^ mice were irradiated and reconstituted with WT or *Nfe2l2*^-/-^ bone marrow cells. Mice were fed a high fat diet for 4 or 10 weeks. Made with BioRender. (**B**-**E**) Aortic roots sections of mice in (**A**) were stained with oil red O (**B, C**) and hematoxylin-eosin (H&E) (**C**). The lesion area and necrotic core are respectively highlighted with a red and yellow dotted line and were quantified (**D, E**). The necrotic core is expressed as percentage of total lesion. Bar graphs show means ± SD and statistical significance was calculated using t-tests.

To investigate which cells respond to oxPAPC activating NRF2 in the atherosclerotic plaque, we analyzed 16,422 myeloid cells from 11 publicly available scRNAseq datasets on aortic immune cells of mice with atherosclerosis^45-50^ (**Figures 5A and 5B**). We identified two cell populations corresponding to monocyte-derived macrophages and *Gpnmb*^+^ foamy macrophages, which were enriched for both oxPAPC and NRF2 signatures (**Figures 5C – E**). Of note, among these cells, only monocyte-derived macrophages (mo-macs) (*Plac8*, *Ear2*, *S100a8, Adgre1, Fcgr4*) expressed high levels of *Il1b* (**Figures 5E**). The monocyte-derived macrophages also expressed high levels of the NRF2-target genes *Gsr*, *Aldoa* and *Taldo1*. These scRNAseq datasets demonstrate that within the plaque, a population of monocyte-derived macrophages respond to oxPAPC, activate NRF2-dependent programs, and produce *Il1b*.

**Figure 5:**
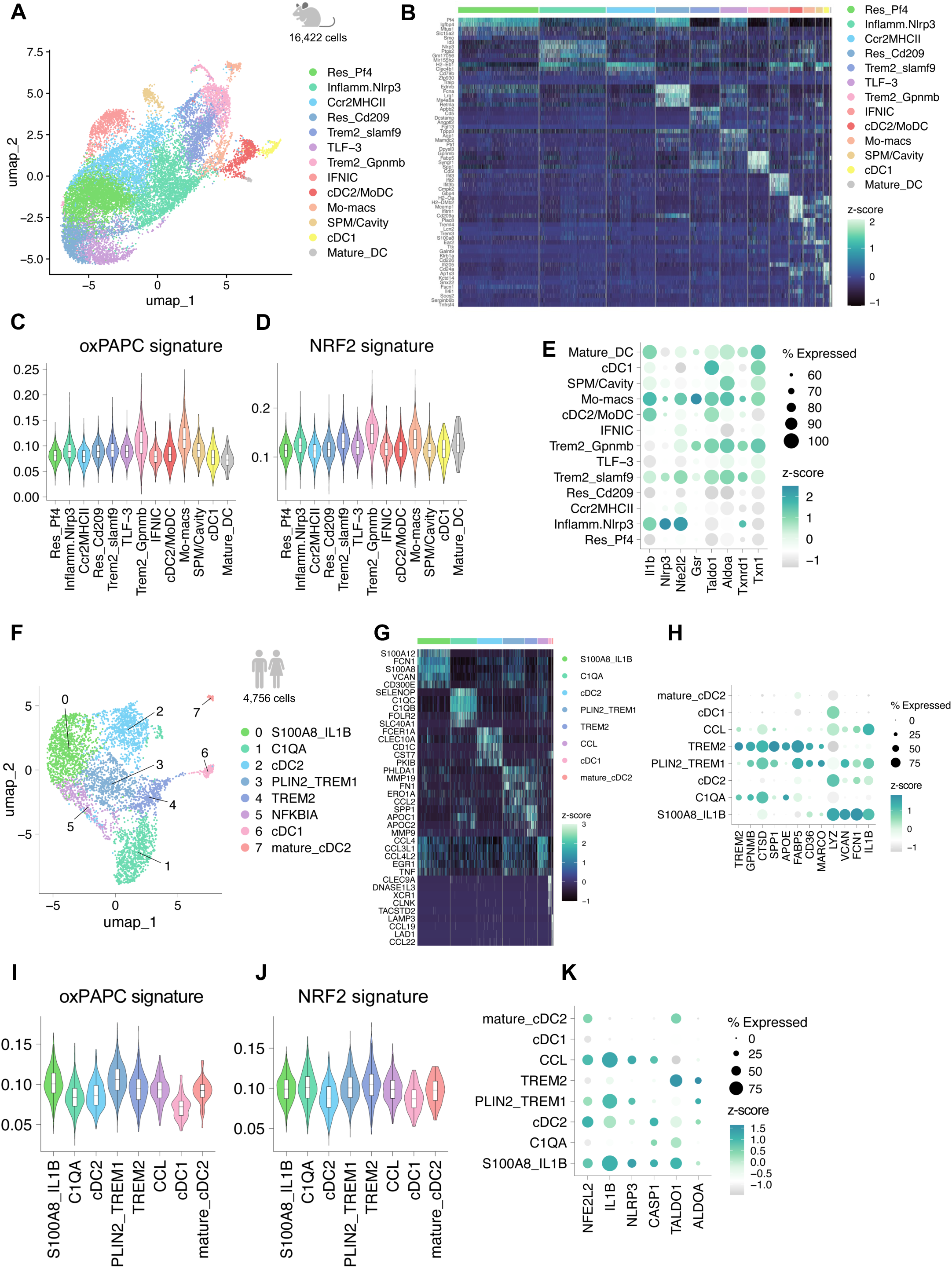
NRF2-dependent signaling characterizes myeloid cells in the mouse and human atherosclerotic plaque that express *NLRP3* and produce *IL1B*. (**A-E**) Integrative scRNAseq analysis of murine aortic immune cells, 10 datasets were reanalyzed from GSE116240, GSE154921, GSE154817, GSE97310, GSE123587, GSE135310^45-50^. UMAP plot of 16,422 cells distributed in the 13 myeloid clusters identified (**A**). Heatmap of the top 5 DEGs used for clustering (**B**). Violin plot representing the gene set scoring of indicated signature across the myeloid clusters identified (**C, D**). oxPAPC signature corresponds to the expression of the top 100 genes upregulated at 3 h and 24 h by oxPAPC (Fig. 2B, C) and the NRF2 signature represents the signature of NRF2-target genes (mouse orthologs of WP2376^70^). (**E**) Dot plot representing the indicated gene expression across myeloid clusters. (**F**-**K**) scRNAseq analysis of immune cells isolated from human atheroma where reanalyzed from GSE210152^51^. UMAP plot of 4,756 myeloid cells, clustered in 8 populations (**F**). Heatmap of the top 5 differentially expressed genes (DEG) used for clustering (**G**). Dot plot representing the indicated gene expression across the myeloid clusters identified (**H**). Violins plot representing the gene set scoring for the human orthologs of the top 100 genes upregulated by oxPAPC at 3 h and 24 h (**I**) and NRF2-regulated genes^71^ (**J**). Dot plot representing the indicated gene expression across the myeloid clusters identified (**K**).

We, next, assessed whether oxPAPC regulates similar programs in the human atherosclerotic plaque. Normalized enrichment score (NES) analysis of bulk RNAseq performed on human atheroma or adjacent intact tissues revealed that the human orthologs of the top 250 genes upregulated by oxPAPC in mouse macrophages (**Figures 2B – D**), as well as the NRF2 signature, were significantly enriched in the atheroma compared to healthy tissue (**Figure S4A**). Next, we analyzed the oxPAPC and NRF2 signatures in 4,756 human myeloid cells derived from the carotid plaque of patients and analyzed by scRNAseq^51^. We identified 8 different functional subsets of myeloid cells (**Figures 5F and 5G**) comprising lipid associated macrophages (C1QA^+^, PLIN2^hi^/TREM1^hi^ and TREM2^hi^) expressing genes linked to lipid uptake and metabolism (*TREM2*, *GPNMB*, *SPP1*, *FABP4*, *CD36*, *MARCO*), S100A8^+^/IL1B^+^ monocytes (*S100A8*, *LYZ*, *IL1B*, *VCAN, CD300E*), inflammatory CCL^hi^ macrophages (*CCL4*, *CCL3L1, CCL4L2, TNF*), and dendritic cells (*CLEC10A*, *CD1C*, *CLEC9A*, *DNASE1L3*) (**Figures 5G and 5H**). The oxPAPC signature was enriched in the S100A8^+^/IL1B^+^ and PLIN2^hi^/TREM1^hi^ clusters (**Figure 5I**), while the NRF2 signature was elevated in several clusters corresponding to monocytes and lipid-associated macrophages (S100A8^+^/IL1B^+^, C1QA^+^, PLIN2^hi^/TREM1^hi^ and TREM2^hi^) (**Figure 5J**). To establish which cells acquired the capacity to secrete mature IL-1β in response to oxPAPC-driven NRF2 and NLRP3 activation, we examined the expression of *IL1B* and *NLRP3* in the different clusters previously identified, and found that 3 clusters displayed high expression of both genes: S100A8^+^/IL1B^+^ monocytes, PLIN2^hi^/TREM1^hi^ inflammatory lipid-associated macrophages, and CCL^hi^ macrophages (**Figure 5K**). Overall, the analysis of the human atherosclerotic plaque confirmed the existence of a monocytic population that responds to oxPAPC by upregulating NRF2-dependent programs and that expresses both *IL1B* and *NLRP3*. It also revealed the presence of a PLIN2^hi^/TREM1^hi^ macrophage population that, despite being lipid-associated, has been described to be proinflammatory and to derive from TREM2^hi^ macrophages^51^. This population responds to oxPAPC by activating a NRF2 transcriptional program and also expresses *IL1B*.

Taken together, these data demonstrate that in the atherosclerotic plaque of both mice and humans it exists a population monocytes and/or macrophages that is characterized by the gene signatures of oxPAPC and NRF2 and that can serve as the local source of bioactive IL-1β, sustaining atherogenesis.

## Discussion

IL-1β is a major driver of metaflammation and its production by macrophages residing in the intima sustains atherosclerosis development^21^. Nevertheless, the nature of triggers and pathways that lead to inflammasome activation in aortic myeloid cells during atherogenesis is still unclear. Here, we focused on oxPLs that are increased in individuals at risk for atherosclerosis development^20, 52^ and that accumulate in the atherosclerotic plaques^20^. We show that oxPLs activate NRF2 that provides both the priming and activation signals to drive the assembly and activity of a functional NLRP3 inflammasome via an all-in-one process that does not require priming with PAMPs. The identification of a host-derived DAMP that simultaneously provides both signals necessary for the activity of NLRP3 is fundamental to explain the contribution of the NLRP3 inflammasome in the context of non-communicable metabolic diseases that lack any overt infection. Mechanistically, we demonstrate that oxPAPC activates the antioxidant transcription factor NRF2. oxPAPC-dependent NRF2 activation on the one hand primes NLRP3 by a yet-to-be defined mechanism and, on the other hand, provides the activation signal for the NLRP3 inflammasome via the production of mtROS and ox-mtDNA. Overall, we demonstrate that not only NRF2 activation by oxPAPC is necessary, but it is also sufficient to drive both the priming and activation of NLRP3.

In mice, plaque formation begins with the uptake of lipids by intimal resident macrophages, followed by a wave of recruited monocytes that play a crucial role in driving plaque progression^45,53^. Our chimeras revealed that NRF2 activation in the hematopoietic compartment is fundamental to drive formation of the atherosclerotic plaque. Re-analyses of scRNAseq dataset also demonstrate that monocyte-derived macrophages, and/or macrophages, are the primary cells within the atherosclerotic plaque that respond to oxPLs activating NRF2 and that are the major cells expressing *NLRP3* and *IL1B*.

Notably, NRF2 is well-known to regulate a potent cytoprotective and anti-inflammatory transcriptional network and its activation is a target of numerous clinical trials against inflammatory diseases^54^. In keeping with its anti-inflammatory role, upregulation of antioxidant programs in macrophages exposed to oxPLs *in vitro* or *in vivo* has been shown to be, at least partially, anti-inflammatory^55-58^. NRF2 activation is generically believed to be protective in metabolic diseases and mouse models of atherosclerosis^54, 59-64^. Nevertheless, scattered reports suggested NRF2 can also play a pathological role^38, 65^, although interpretation of the relevance of NRF2-signaling in immune cells is confounded by the fact that, in these investigations, both stromal and immune cells lack NRF2. Our *in vitro* and *in vivo* data clearly establish a direct link between NRF2 activation and NLRP3 priming and activation. Although a previous study found that NRF2 is involved in driving inflammasome activation in response to Nigericin^40^, our and previous data^41^ with ATP and Nigericin ruled out the involvement of NRF2 in the activation of the NLRP3 inflammasome in response to canonical NLRP3-activating stimuli. These discrepancies may be due to specific cell culture conditions that can impact distinct parameters, such as cellular density and/or the concentration of the NLRP3 agonist utilized.

Our investigation also established that NRF2 drives the activation of NLRP3 via the formation ox-mtDNA. Regarding the priming of NLRP3, our data demonstrate that NRF2 prolonged activation is sufficient to trigger inflammasome activity in the absence of PAMP detection. During the priming phase, NLRP3 and, to a lesser extent, ASC and caspase-1 have been shown to be regulated by a wide variety of PTMs, including phosphorylation, ubiquitination, and palmitoylation^7, 66-69^. Further investigation will determine how oxPAPC-mediated NRF2 activation primes NLRP3, and whether this priming occurs through posttranslational modifications of NLRP3.

In conclusion, we demonstrate that oxPLs act as all-in-one NLRP3 inflammasome primers and activators through the manipulation of the cellular redox of monocytes and macrophages, driving the production of bioactive IL-1β and participating to the pathogenesis of atherosclerosis in mice and humans.

## Acknowledgment

IZ was supported for these studies by NIH grant R01AI170632, and Lloyd J. Old STAR Program CRI3888. RS thanks the UCLA Institute for Quantitative and Computational Biosciences (QCBio) Collaboratory community directed by Matteo Pellegrini. OD received a mobility grant from the Philippe Foundation. M.D.G. is supported by the Crohn’s and Colitis Foundation, award number 854391. K.W. is supported by a NIH-NIAMS K08AR077037, a Burroughs Wellcome Fund Career Awards for Medical Scientists, and a Doris Duke Charitable Foundation Clinical Scientist Development Award.

## Author contribution

OD performed *in vitro* and *in vivo* experiments, conceived and analyzed the experiments, and wrote the manuscript; MDG performed the bulk RNA-seq experiments and helped to analyze data and to conceive experiments; RS performed the bioinformatics analyses; PJT and JRS prepared oxPAPC; SAP, JC, CG, KW helped to analyze scRNAseq data; HW provided the human monocytic cell line and gave advice; IZ conceived and supervised the experiments, and wrote the paper.

## Data availability

All transcriptional data will be provided upon request to the corresponding author.

---

**Figure S1.**
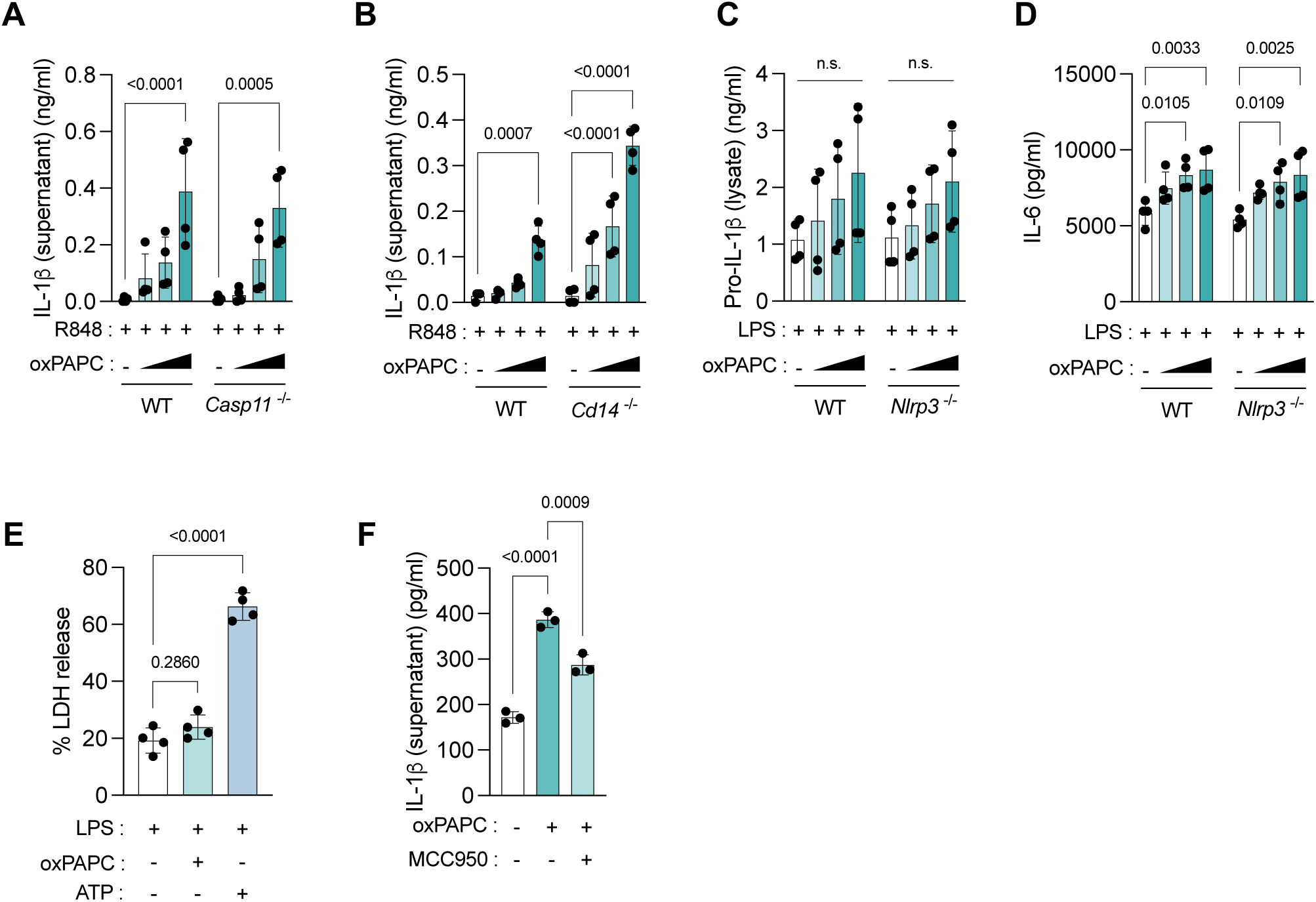
(**A**-**D**) WT, *Casp11*^-/-^, *Cd14*^-/-^ or *Nlrp3*^-/-^ BMDMs were pre-conditioned with oxPAPC (100 µg/ml) for 24 h and then stimulated with R848 or LPS (1 µg/ml) for 24 h. Supernatants (**A, B, D**) or cell lysates (**C**) were analyzed by ELISA. (**E**) BMDMs were either pre-conditioned with oxPAPC (100 µg/ml) for 24 h and then stimulated with LPS (1 µg/ml) for 24 h or treated for 4 h with LPS (1 µg/ml) before adding ATP (5 mM) for 1 h. Lactate dehydrogenase (LDH) activity was assessed in the supernatant. (**F**) THP-1 monocytes expressing mTurquoise2-hProIl1b-mNeonGreen were treated with MCC950 (5 µM) for 1 h before adding oxPAPC (100 µg/ml) for 24 h. Supernatants were analyzed by ELISA. Bar graphs show means ± SD and statistical significance was calculated using a Two-way ANOVA.

**Figure S2.**
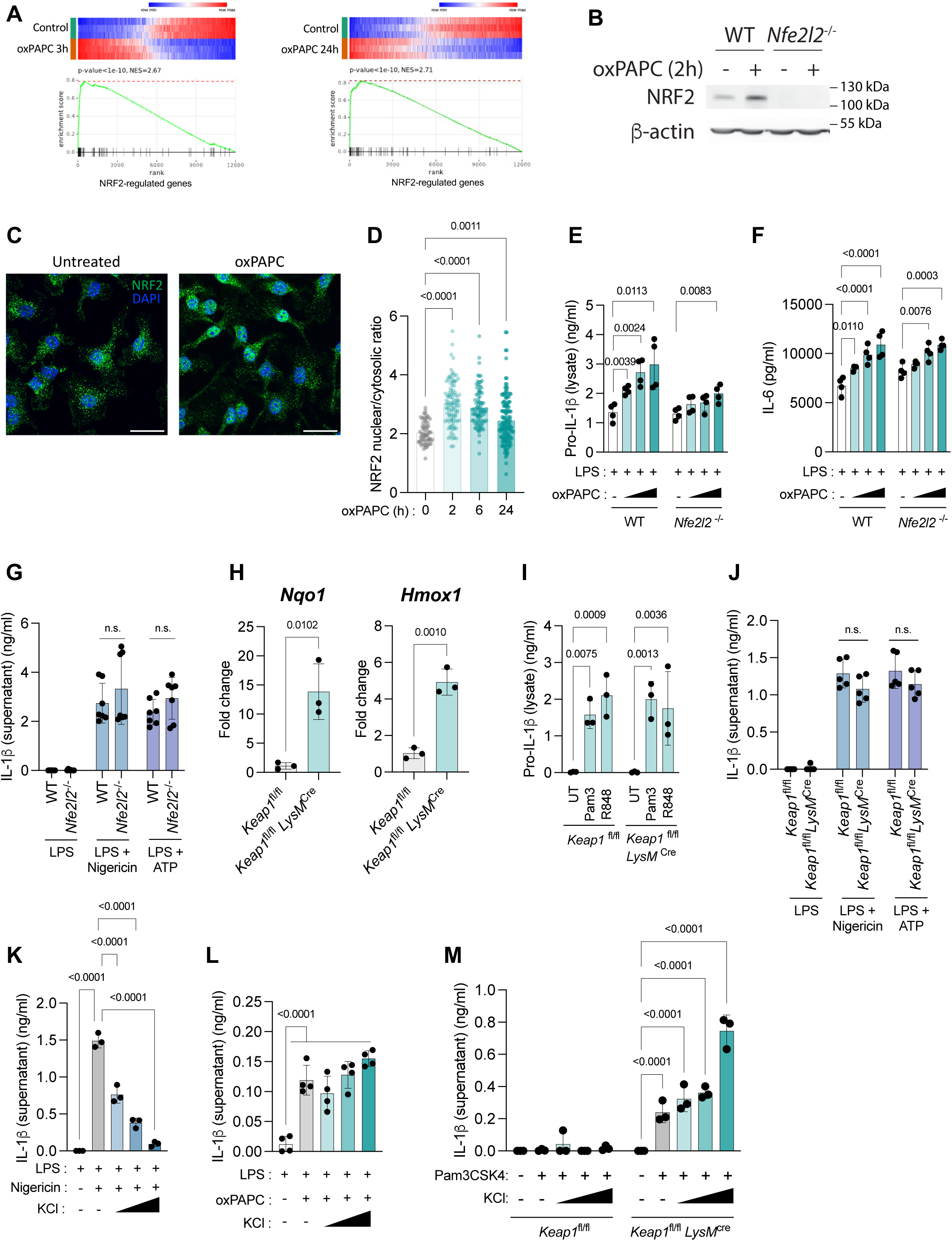
(**A**) Gene set enrichment analysis (GSEA) plot of NRF2-regulated genes (mouse orthologs of WP2376^70^) comparing the transcriptional profile of BMDMs conditioned with oxPAPC for 3 h or 24 h and BMDMs left untreated. The graph shows the enrichment score (ES) for NRF2-regulated genes. (**B**-**D**) BMDMs were conditioned with oxPAPC for 2 h (**B**-**D**), 6 h or 24 h (**D**). Whole cell lysates were analyzed by immunoblot (**B**). BMDMs were stained with an anti-NRF2 antibody (green) and DAPI (blue, nuclei) (**C**), and the nuclear-cytoplasmic ratio of NRF2 distribution was quantified (**D**). (**E, F**) WT and *Nfe2l2*^-/-^ BMDMs were conditioned with oxPAPC (100 µg/ml) for 24 h and treated with LPS (1 µg/ml) for 24 h. Cell lysates (**E**) and supernatants (**F**) were analyzed by ELISA. (**G**) WT and *Nfe2l2*^-/-^ BMDMs were primed for 4 h with LPS (1 µg/ml) before treatment for 1 h with Nigericin (5 µM) or ATP (5 mM). Supernatants were analyzed by ELISA. (**H**) Untreated *Keap1*^fl/fl^ and *Keap1*^fl/fl^*LysM*^cre^ BMDMs were analyzed for *Nqo1* and *Hmox1* expression by reverse transcriptase-qPCR. (**I**) *Keap1*^fl/fl^ and *Keap1*^fl/fl^*LysM*^cre^ BMDMs were treated with R848 or Pam3 (1µg/ml) for 24 h. Cell lysates were analyzed by ELISA. (**J**) *Keap1*^fl/fl^ and *Keap1*^fl/fl^*LysM*^cre^ BMDMs were treated with LPS for 4 h before adding Nigericin (5 µM) or ATP (5 mM) for 1h. Supernatants were analyzed by ELISA. (**K**) WT BMDMs were treated as in (**J**) in a medium containing 10, 20 or 30 mM KCl. Supernatants were analyzed by ELISA. (**L**) WT BMDMs were treated as in (**E**) in a medium containing 10, 20 or 30 mM KCl. Supernatants were analyzed by ELISA. (**M**) *Keap1*^fl/fl^ or *Keap1*^fl/fl^*LysM*^cre^ BMDMs were treated as in (**I**) in a medium containing 10, 20 or 30 mM KCl. Supernatants were analyzed by ELISA. Bar graphs show means ± SD and statistical significance was calculated using a Two-way ANOVA and t-tests.

**Figure S3.**
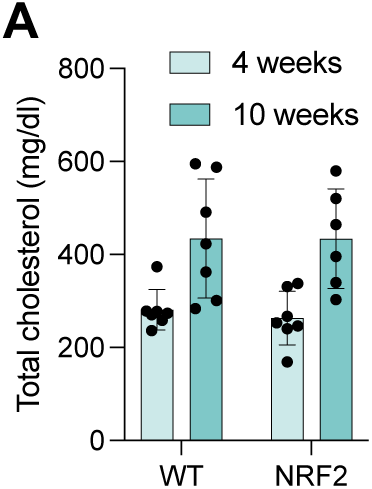
(**A**) Total cholesterol measurement from plasma of mice showed in Figure 4A-E.

**Figure S4.**
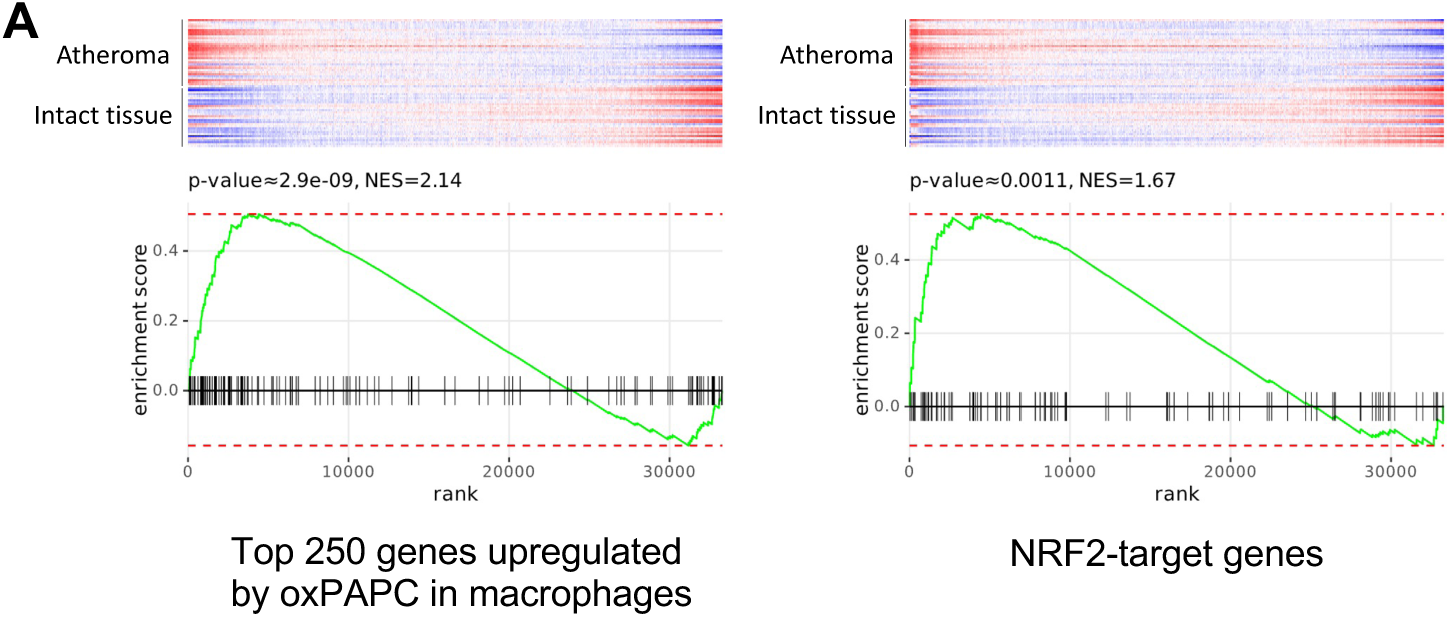
(**A**) GSEA plot of the human orthologs of the top 250 genes upregulated by oxPAPC in BMDMs (*left*) and NRF2-target genes (*right*) comparing the transcriptional profile of human atheroma and healthy tissue of the same patient (GSE43292^51^).

## Methods

### Mice

The animals were housed at the Boston Children’s Hospital animal facility under controlled temperature, and with a 12:12 h light-dark cycle. All animals had *ad libitum* access to chow diet and water. All animal experiments were performed under the Guide of Care and Use of Laboratory Animals of the National Institute of Health and all animal experiment were approved by the Institutional Animal Care and Use Committee (IACUC) of Boston Children’s Hospital. The mouse strains used in this study were purchased at the Jackson Laboratory and were the following: C57BL/6J (WT, Jax 000664), B6.Cg-*Casp1*^em1Vnce^/J (*Casp1*^-/-^, Jax 032662), B6.129S6-*Nlrp3^tm1Bhk^*/J (*Nlrp3*^-/-^, Jax 021302), B6.129S4(D2)-*Casp4^tm1Yuan^*/J (*Casp11*^-/-^, Jax 024698), B6.129S4-*Cd14^tm1Frm^*/J (*Cd14*^-/-^, Jax 003726), B6.129X1-*Nfe2l2^tm1Ywk^*/J (*Nfe2l2*^-/-^, Jax 017009), B6(Cg)-*Keap1*^tm1.1Sbis/^J (*Keap1*^fl/fl^, Jax 037075), B6.129P2-*Lyz2^tm1(cre)Ifo^*/J (*LysM*^cre^, Jax 004781), B6.129S7-Ldlrtm1Her/J (*Ldlr*^-/-^, 002207). *Keap1*^fl/fl^*LysM*^cre^ were obtained by crossing *Keap1*^fl/fl^ and *LysM*^cre^ mice.

### Murine model of atherosclerosis and bone marrow chimeras

Sulfatrim (Trimethoprim / Sulfamethoxazole) was added to the drinking water of the recipient mice 1 week before irradiation and for 2 weeks after bone marrow reconstitution. 8 weeks-old *Ldlr*^-/-^ female mice were lethally irradiated with a total dose of 950 rad (γ-irradiation, cesium^137^-based) in a Gammacell 40 Exactor (Best Theratronics). Mice first received 475 rad and after a resting time of 3 hours, the irradiation was repeated (475 rad). 4 hours after the last irradiation, mice received freshly isolated bone marrow from female, 6 to 8 weeks old, WT or *Nfe2l2*^-/-^ donor mice as previously described^72^. After isoflurane-based anesthesia, mice received 3×10^6^ bone marrow cells in 200μl of RPMI 1640 via retro-orbital injection. Mice were kept in autoclaved cages for 2 weeks after reconstitution. After a resting period of 4 weeks, mice were fed a high fat diet (HFD) containing 42% kcal, 0.2% cholesterol (TD.88137, Teklad). After 4 or 10 weeks of HFD, mice were euthanatized with 100% CO_2_ and infused with 20 ml of ice-cold PBS. Hearts were collected and embedded in OCT and stored at -80 °C.

### Aortic root staining and quantification

10 µm serial sections of the aortic root were stained with oil red O and hematoxylin-eosin (H&E) by iHisto (Salem, MA, USA). The lesion was defined as positive for oil red O staining. The necrotic core was defined using H&E and as region with low cellularity. Necrotic core area is expressed as the percentage of the total lesion. Sections were analyzed with Fiji version 2.14.0 (ImageJ).

### Bone marrow-derived macrophages and THP-1 monocytes culture

Mice were euthanized with 100% CO_2_ and bone-marrow was collected from femurs and tibias by flushing the bones with PBS. Bone marrow cells were differentiated into macrophages at 37 °C, 5% CO_2_ for 6-7 days in DMEM (Thermo Fisher Scientific, Cat# 11965092) containing 30% L-929 conditioned media and supplemented with 10 % heat-inactivated FBS (Thermo Fisher Scientific, Cat# A5256701), 2 mM L-glutamine (Thermo Fisher Scientific, Cat# 25030081) and 100 μg/ml penicillin/streptomycin (Thermo Fisher Scientific, Cat# 15140122). THP-1 monocytes stably expressing mTurquoise2-IL-1b-mNeongreen^30^ were maintained in RPMI 1640 GlutaMAX (Thermo Fisher Scientific, Cat# 61870036) containing 10% heat-inactivated FBS (Thermo Fisher Scientific, Cat# A3382001) and 100 μg/ml penicillin/streptomycin (Thermo Fisher Scientific, Cat# 15140122).

### Cell treatment

BMDMs were cultured in DMEM (Thermo Fisher Scientific, Cat# 11965092) containing 10% heat-inactivated FBS (Thermo Fisher Scientific, Cat# A5256701), 2 mM L-glutamine (Thermo Fisher Scientific, Cat# 25030081) and 100 μg/ml penicillin/streptomycin (Thermo Fisher Scientific, Cat# 15140122). Macrophages were incubated as indicated in the figure legends with oxPAPC (25, 50 or 100 µg/ml), ultrapure LPS from *E. coli* 055:B5 (Invivogen, Cat# tlrl-pb5lps) 1 µg/ml, R848 (Invivogen, Cat# tlrl-r848-1) 1 µg/ml, Pam3CSK4 (Invivogen, Cat# tlrl-pms) 1 µg/ml, Nigericin 5 µM (Invivogen, Cat# tlrl-nig) or ATP (pH 7.4) 5mM (Sigma-Aldrich Cat# A6419). When inhibitors were used, cells were pre-treated for 1 hour with the following inhibitors (the concentrations used are indicated in the figure legends): MCC950 (MedChemExpress, Cat# HY-12815), Mitoquinol (MedChemExpress, Cat# HY-124410), FEN1-IN-4 (MedChemExpress, Cat# HY-136485), VBIT-4 (MedChemExpress, Cat# HY-129122). Sterile-filtered KCl (Sigma-Aldrich, Cat# P5405) was added to the media concurrently to the NLRP3-activating stimuli (oxPAPC or Nigericin).

### ELISA

Cells were cultured in 96-well plates and the plate was centrifuged for 5 min at 1500 rpm. Supernatant was collected and cell lysed with PBS containing 1% Triton X-100 and protease/phosphatase inhibitor (Thermo Fisher Scientific, Cat# 1861284). ELISA were performed according to the manufacturer instruction using ELISA MAX Standard Set Mouse for IL-1β (Biolegend, Cat# 432601) or IL-6 (Biolegend, Cat# 431301) or ELISA MAX Deluxe Set Human IL-1β (Biolegend, Cat# 437004). Briefly, ImmunoGrade 96-well plates (Genesee, Cat# 91-415F) were coated with the capture antibody overnight at 4 °C, washed 3 times with PBS 0.05% Tween-20 and blocked for 1 hour at room temperature with the assay diluent A from the kit. After 1 wash with PBS 0.05% Tween-20, samples and standards for hIL-1β, mIL-1β or mIL-6 were incubated overnight. The plate was washed 3 times with PBS 0.05% Tween-20 and the appropriate detection antibody was added for 1 hour, at room temperature. After 3 washes, Avidin-HRP was added for 30 min at room temperature and in the dark. Plates were washed 5 times with PBS 0.05% Tween-20 before adding the substrate solution D and stopped with 2N H_2_SO_4_ solution. ELISA plates were read on the SpectraMax i3x plate reader (Molecular Devices).

### Western blot and immunostaining

At the end of the treatment, supernatants were collected and Laemmli 4X (Bio-Rad, Cat# 1610747) and DL-Dithiothreitol (Sigma-Aldrich, Cat# D0632) was added to a final concentration of 50mM. Cells were lysed in Laemmli 2X supplemented with 50mM DL-Dithiothreitol (Sigma-Aldrich, Cat# D0632). Sample were boiled and loaded into 10 to 15% acrylamide gels. After transfer and blocking in 5% non-fat dry milk, membrane were probed with the following antibodies: CASP1 p20 (Adipogen, Cat# AG-20B-0042), IL-1β (Genetex, Cat# GTX4034), NLRP3 (Abcam, Cat# ab270449), β-ACTIN (Sigma–Aldrich, Cat# A5441), KEAP1 (Cell Signaling, Cat# 8047S), NRF2 (Cell Signaling, Cat# 12721S), GAPDH (Cell Signaling, Cat# 97166), Histone H3 (Cell Signaling, Cat# 4499S).

For immunofluorescence, cells were plated on coverslips and after treatment were washed once with PBS and were fixed with PBS 4% paraformaldehyde (Thermo Fisher Scientific, Cat# 047377.9M) for 15 min at room temperature. Paraformaldehyde was quenched with 50mM NH_4_Cl for 15 min and cells were permeabilized with PBS 0.5%Triton for 5 min. Cells were incubated for 1 hour with TBS, 5% BSA, 0.1% Tween-20 at room temperature to block unspecific signal. The primary antibodies were used 1:100 in TBS, 5% BSA, 0.1% Tween-20 and the staining was performed overnight at 4 °C in a humidity chamber. The next day, cells were stained with secondary antibodies (1:400) and DAPI (300nM) (Thermo Fisher Scientific, Cat# D1306) for 1 hour, at room temperature. The coverslips were mounted on slides with Prolong gold antifade reagent (Thermo Fisher Scientific, Cat# P36934). The following antibodies were used for immunofluorescence: NRF2 (Cell Signaling, Cat# 12721S), 8-OHdG (Abcam, Cat# ab62623) and TOM20 (Abcam, Cat# ab186735), Goat anti-Rabbit IgG Alexa Fluor 568 (Thermo Fisher Scientific, Cat# A11011), Chicken anti-Mouse IgG Alexa Fluor 488 (Thermo Fisher Scientific, Cat# A21200).

Immunofluorescence were analyzed with a Zeiss LSM 880 microscope using the Airyscan mode. Images were analyzed with Fiji version 2.14.0 (ImageJ).

### Cell fractionation

5×10^6^ BMDMs were treated as indicated and washed once with cold PBS (Genesee, Cat# 25-508). Cells were lysed in subcellular fractionation buffer (20 mM HEPES pH 7.4, 10 mM KCl, 2 mM MgCl_2_, 1 mM EDTA, 1mM EGTA, 1mM DL-Dithiothreitol and protease/phosphatase inhibitor (Thermo Fisher Scientific, Cat# 1861284)) and lysates were incubated on ice for 15 min. Lysates were passed 15 times through a 27 gauges needle and were incubated on ice for 20 min, then centrifuged at 720 x g for 5 min at 4 °C. The pellet was resuspended with 500 µL of fractionation buffer by passing 10 times through a 27 gauges needle and centrifuged at 720 x g for 10 min at 4 °C. The pellet was resuspended in Laemmli buffer containing 50 mM DL-Dithiothreitol (Sigma-Aldrich, Cat# D0632). The supernatant from the first centrifugation was kept and centrifuged at 10,000 x g for 10 min at 4 °C before adding Laemmli buffer and 50 mM DL-Dithiothreitol. Nuclear and cytoplasmic fractions were boiled at 95 °C for 20 and 10 min, respectively. Samples were analyzed by western blot.

### LDH release

The supernatant of stimulated macrophages was collected and centrifuged at 300g for 5 min to remove cellular debris. LDH measurement was performed using the CyQUANT LDH Cytotoxicity Assay Kit (Thermo Fisher Scientific, Cat# C20300) according to the manufacturer’s instructions. Data are shown as the percentage of LDH release considering a Triton X-100 treated well as 100%.

### Quantitative RT-PCR

RNA was extracted with Direct-zol RNA Miniprep Kit (Zymo Research, Cat# R2053) and purified RNA was analyzed using Power SYBR Green RNA-to-CT 1-Step Kit (Thermo Fisher Scientific, Cat# 4389986) with predesigned primers (Sigma-Aldrich, KiCqStart SYBR Green Primers) specific to murine *Hmox1*, *Nqo1* and *Rpl13a*.

### Mitochondrial ROS and flow cytometry

Mitochondrial ROS were stained with mitoSOX red (Invitrogen, Cat# M36008) as per manufacturer instructions. Briefly, after treatment, cells were stained in HBSS media containing 5 µM of freshly reconstituted MitoSOX red and were incubated at 37 °C for 30 min. Cells were washed once in PBS and analyzed by flow cytometry with a LSRFortessa (Becton, Dickinson and Company). Data were then analyzed using FlowJo v10.10 (Becton, Dickinson and Company).

### Bulk RNAseq and analysis

1×10^6^ BMDMs were treated left untreated or treated for 3 hours or 24 hours with oxPAPC. RNA was purified using Direct-zol RNA Miniprep (Zymo Research Corporation, Cat# R2053). Samples were submitted to Azenta Life Sciences for standard RNA-seq processing. RNA was quantified using Qubit 2.0 Fluorometer (Thermo Fisher Scientific) and RNA integrity was checked with RNA Screen Tape on Agilent 2200 TapeStation (Agilent Technologies). After library preparation (Illumina, PolyA selection), samples were sequenced using Illumina, 2×150 bp, ∼350M PE reads (∼105GB), single index. Reads were quality-controlled using FastQC. Illumina adapters were removed using cutadapt. Trimmed reads were mapped to the mouse transcriptome (GRCm38) based on Ensembl annotations using Kallisto. Gene counts and differential expression were analyzed using the R package Limma^73^. GSEA enrichment plots and FGSEA were performed using the online application Phantasus^74^.

### Human dataset scRNAseq analysis

The scRNAseq dataset of immune cells isolated from human atheroma was reanalyzed from GSE210152^51^ using the Seurat^75^ R package (version 5.1.0). Cells were filtered to retain those with more than 500 detected genes and less than 5% mitochondrial gene content. The filtered dataset was then normalized using SCTransform, and highly variable features were identified using the VST method, with the top 2000 variable features selected and PCA was performed on the identified variable features. Samples were integrated using the Harmony algorithm^76^. The nearest neighbors graph was created using the first 30 principal components, and clustering was performed using the Louvain algorithm with a resolution parameter of 0.3. UMAP was applied to visualize the clusters. Non-myeloid clusters were identified and removed, and the remaining myeloid subset was analyzed as before. This included SCTransform, identification of variable features, PCA, integration and construction of a nearest neighbors graph, clustering with a resolution parameter of 0.4, and UMAP visualization. Marker genes for each cluster were identified, selecting genes expressed in more than 30% of cells with at least a 0.3 log fold change. We identified 8 populations with a total of 4,756 myeloid cells. Assignation of each cell type was done according to the initial study. The gene set scoring for oxPAPC signature (human orthologs of the top 100 genes upregulated by oxPAPC at 3 and 24 hours in macrophages) and NRF2-regulated genes (WP2884^71^) was perform using the UCell package^77^ on R.

### Murine datasets scRNAseq analysis

We reanalyzed 10 scRNAseq publicly available datasets from 5 studies (GSE116240, GSE154921, GSE154817, GSE97310, GSE123587, GSE135310)^45, 46, 48-50^ following the workflow used to integrate the datasets in Zernercke et al.^47^ using the Seurat package^75^ (version 5.1.0) on R. Each scRNAseq dataset was individually pre-processed as described in Zernercke et al.^47^. Cells were filtered to include those with more than 200 detected genes, fewer than 25,000 RNA counts, and less than 5% mitochondrial genes. The datasets were then normalized, and highly variable features were identified using the VST method, with the top 2000 variable features selected. Data were scaled prior to performing PCA on the identified variable features. The nearest neighbors graph was constructed using 14 or 20 PCs as determined by the JackStraw analysis. Clusters were then identified using the Louvain algorithm with a resolution of 0.8 on 14 to 20 PCs. The datasets were then merged into a unique Seurat object for integrated analysis. Normalization and identification of highly variable features were performed for each dataset, and integration anchors were identified using 30 PCs. The integrated dataset was scaled and PCA was conducted, with the first 30 PCs used for further analysis. UMAP was then applied based on the first 30 PCs. An SNN graph was constructed using 30 PCs, and clustering was performed using the Louvain algorithm with a resolution parameter of 0.6. Marker genes for each cluster were identified. For each cluster, genes that were expressed in more than 30% of cells with at least a 0.3 log fold change were selected. Non-myeloid clusters were identified and removed, and cells that segregated with the non-myeloid clusters on the UMAP coordinates were manually removed. We identified 16,422 cells distributed in 13 myeloid clusters identified, the annotation of each cluster was made according to the marker genes and based on Zernercke et al.^47^. The gene set scoring for oxPAPC signature (human orthologs of the top 100 genes upregulated by oxPAPC at 3 and 24 hours in macrophages) and NRF2-regulated genes (WP2376^70^) was perform using the UCell package^77^.

### oxPAPC and NRF2 signatures

The oxPAPC signature was defined as the top 100 genes upregulated at 3 hours and 24 hours following oxPAPC treatment in BMDMs. After removing duplicates, the final oxPAPC signature included 168 unique genes. The NRF2 signature was defined as the mouse orthologs of NRF2-regulated genes curated from WikiPathways WP2376^70^. For human, the NRF2 signature was curated from WikiPathways WP2884^71^.

### PAPC oxidation

PAPC was oxidized as previously reported^25, 78, 79^. Briefly, PAPC (Avanti Lipids) was transferred to clean borosilicate tubes in 1 mg aliquots in chloroform, dried, and oxidized for 24–72 hours, while monitoring oxidation by flow injection on an electrospray ionization (ESI) instrument (Thermo LCQ). Lipids were analyzed by phosphorus assay to determine the concentration.

### Statistical analysis

Statistical significance for the experiments with more than two groups and two factors was tested with two-way ANOVA, and Sidak’s, Dunnet’s or Tukey’s multiple-comparisons tests were performed according to the nature of the comparison tested. When comparisons between only two groups were made, an unpaired two-tailed t-test was used to assess statistical significance. Except for bulk RNAseq analyses, all statistical analyses were performed using Prism 9 (GraphPad Software, version 10.2.1). For bulk RNA-seq analysis, the adjusted p-value was calculated with the limma package^73^ on R.

